# Repression by hdac3 and dax1 mediates lineage restriction of embryonic stem cells

**DOI:** 10.1101/2020.09.10.291013

**Authors:** Daniel Olivieri, Panagiotis Papasaikas, Ilya Lukonin, Melanie Rittirsch, Daniel Hess, Sébastien A. Smallwood, Michael B. Stadler, Joerg Betschinger

## Abstract

Mouse embryonic stem cells (mESCs) give rise to embryonic but not extraembryonic endoderm fates. Here, we identify the mechanism of this lineage barrier and report that the histone deacetylase Hdac3 and the corepressor Dax1 cooperatively restrict transdifferentiation of mESCs by silencing an enhancer of the extraembryonic endoderm-specifying transcription factor (TF) Gata6. This restriction is opposed by the pluripotency TFs Nr5a2 and Esrrb, which promote cell type conversion. Perturbation of the barrier extends mESC potency, and allows formation of 3D spheroids that mimic the spatial segregation of embryonic epiblast and extraembryonic endoderm in early embryos. Overall, this study shows that transcriptional repressors stabilize pluripotency by biasing the equilibrium between embryonic and extraembryonic lineages that is hardwired into the mESC TF network.

## INTRODUCTION

Binary cell fate decisions generate two distinct daughter cell types from one common progenitor. Once specified, cells need to prevent de-differentiation and transdifferentiation in order to stay fate-committed. The processes of lineage specification and maintenance can employ different mechanisms (Holmberg and Perlmann, 2012), yet the regulatory principles underlying these differences are not well understood.

A well-studied developmental binary cell fate decision is the differentiation of the mouse embryonic day (E) 3.5 inner cell mass (ICM) into extraembryonic primitive endoderm (PrE) and pluripotent epiblast (EPI) (Hermitte and Chazaud, 2014). This is mediated by the lineage-specifying transcription factors (TFs) Nanog and Gata6 that are co-expressed in the ICM. Positive auto-regulation and mutual inhibition of Nanog and Gata6, modulated by extrinsic fibroblast growth factor (FGF) signaling, drives segregation into Gata6-expressing PrE and Nanog-expressing EPI at E4.5 (Simon et al., 2018).

Developmental potency of the EPI is captured *in vitro* by mouse embryonic stem cells (mESCs) that are restricted to predominantly give rise to embryonic cell types *in vitro* and when injected into blastocysts (Beddington and Robertson, 1989). The lineage barrier impeding PrE differentiation of mESCs is not well understood. Overexpression of Gata6 enables transdifferentiation into extraembryonic endoderm (XEN) (Shimosato et al., 2007), raising the possibility that the Nanog-Gata6 antagonism that segregates EPI and PrE during mouse pre-implantation development maintains lineage separation in mESCs (Graf and Enver, 2009). Nanog is, however, dispensable for post-implantation development and mESC self-renewal (Chambers et al., 2007), suggesting operation of additional mechanisms that govern fate restriction (Holmberg and Perlmann, 2012), such as the activation of TFs that function redundantly with Nanog.

Different mESC states are stabilized by specific extrinsic signaling conditions (Smith, 2017): A naïve pluripotent groundstate in the presence of 2 inhibitors and leukemia inhibitory factor (LIF) (2iL), and a metastable pluripotent state by foetal calf serum (S) and LIF (SL). Although both states are interconvertible, mESCs in SL heterogeneously express differentiation and pluripotency markers (Smith, 2017), inefficiently differentiate into Epiblast-like cells (EpiLCs) *in vitro* (Hayashi et al., 2011), and are thought to recapitulate a developmentally more advanced state than mESCs in naïve 2iL conditions (Gonzalez et al., 2016; Schröter et al., 2015).

Here, we exploit the transition from the naïve to the metastable mESC state (Schröter et al., 2015) to define the mechanism of lineage restriction. We identify inhibitors of PrE transdifferentiation, define their interaction with pluripotent TFs, and describe how they enact competition between PrE and EPI fate. Our findings reveal that silencing of a Gata6 enhancer by transcriptional repressors antagonizes lineage plasticity of the mESC gene regulatory network (GRN) and secures the pluripotent lineage.

## RESULTS

### Hdac3 inhibits transdifferentiation of mESCs into PrE

While working on a putative Hdac3 interactor, we genetically deleted *Hdac3* in naïve TNG-A mESCs that express GFP under the control of the endogenous *Nanog* locus (Chambers et al., 2007) (**Figure S1A**). Compared to *wildtype* (*WT*) controls, *Hdac3*^**-/-**^ cells expressed higher levels of the Nanog reporter in 2iL, but rapidly downregulated Nanog when converted to SL (**Figure 1A**) and were lost upon further passaging, indicating undue differentiation specifically in metastable conditions.

**Figure 1:**
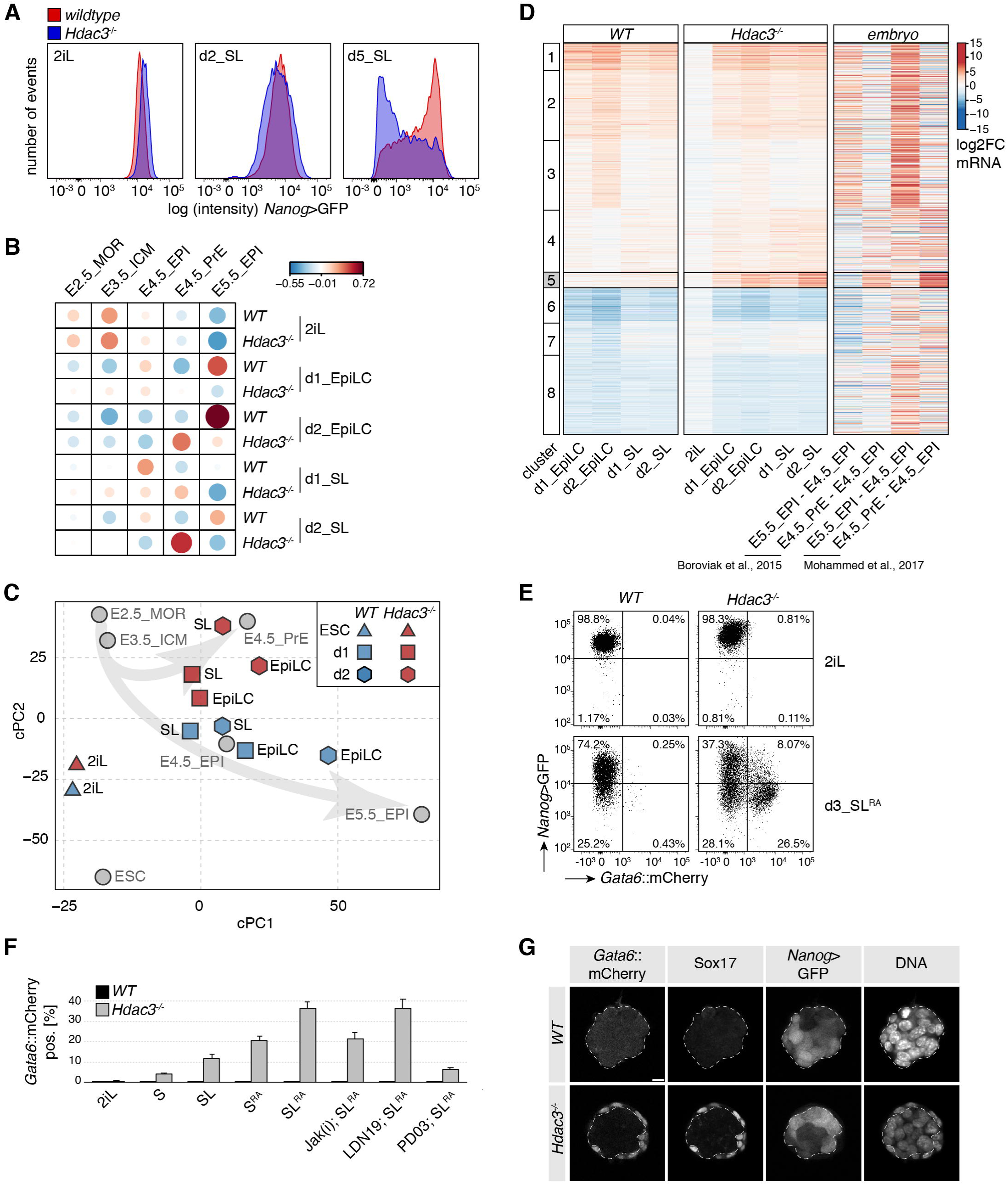
Naïve *Hdac3*^-/-^ mESCs transdifferentiate to PrE in response to FGF, LIF, and RA. **(A)** *Nanog*>GFP intensity of *WT* and *Hdac3*^**-/-**^ TNG-A cells in 2iL and at d2 and d5 of SL exposure. **(B**,**C)** Pairwise Pearson correlations (B) and common PC analysis (C) of *in vitro* TNG-A and *in vivo* embryo (Boroviak et al., 2015) RNAseq samples. Morula (MOR). **(D)** k-means clustering of gene expression changes relative to naïve *WT* TNG-A ESCs. **(E)** *Nanog*>GFP and *Gata6*::mCherry fluorescence intensities of *WT* and *Hdac3*^**-/-**^ cells in indicated conditions. **(F)** Percentage of Gata6::mCherry positive cells after 3d of exposure in indicated conditions. Jak(i) blocks LIF, LDN19 BMP4 and PD03 FGF signaling. Average and SD of three independent clones. **(G)** Immunofluorescence and reporter expression in spheroids derived from single WT and *Hdac*3^**-/-**^ naïve mESCs. DNA counterstain is Hoechst. The embryonic *Gata6*::mRFP negative part is indicated. Scalebar: 10μM.

To determine the transcriptional changes underlying this phenotype, we performed RNA sequencing (RNAseq) of *WT* and *Hdac3*^**-/-**^ cells in the naïve ESC state, and after 1 and 2 days (d) in SL and epiblast-like cell (EpiLC) (Hayashi et al., 2011) differentiation conditions (**Table S2**). Contrasting these results with existing datasets from the early embryo (Boroviak et al., 2015; Mohammed et al., 2017) using pairwise correlation and principal component (PC) analysis (**Figure 1B,C, S1B,C**) revealed that mESCs in 2iL are most similar to the embryonic day E3.5 ICM (Gonzalez et al., 2016). It further showed that SL and EpiLC conditions drive naïve *WT* mESCs into cell states resembling the embryonic E4.5 - E6.5 pre- and post-implantation EPI. In contrast, differentiating *Hdac3*^**-/-**^ cells, in particular in SL, were transcriptionally most similar to primitive and visceral (VE), but not definitive endoderm (Anderson et al., 2017) (**Figure S1D**). Clustering analysis identified a class of genes (cluster 5) that was selectively induced in differentiating mutant cells and the E4.5 PrE *in vivo* (**Figure 1D**). Cluster 5 is enriched for endoderm regulators by gene ontology analysis and includes the TFs Gata4, Gata6, and Sox17 that are required for PrE development *in vivo* (Hermitte and Chazaud, 2014) (**Figure S1E,F**). Absence of *Hdac3* therefore causes PrE transdifferentiation upon transition from naïve to metastable conditions.

Naïve mESCs engineered to express catalytically inactive Hdac3 (*Hdac3*^Y118F/Y118F^), or depleted of *Ncor1* and *Ncor2* by siRNA transfection and compound knockout (Sun et al., 2013) similarly induced PrE markers after SL conversion (**Figure S1G,H, Table S1**), demonstrating that Hdac3 acts as a deacetylase and in Ncor1/2 nuclear corepressor complexes.

### Hdac3 shields mESCs from PrE conversion in response to extrinsic developmental signals

The induction of post-implantation markers in *Hdac3*^**-/-**^ cells was unperturbed (**Figure S1I**), suggesting simultaneous activation of embryonic and extra-embryonic gene expression. To test if this is due to population heterogeneity, we deleted *Hdac3* in a TNG-A-derived ESC line (G6C18) that in addition to *Nanog* reports endogenous transcription of *Gata6* (**Figure S1J**). After 3d in the presence of SL, around 10% of mutant cells were positive for the *Gata6* reporter (**Figure 1E,F, S1K**). Addition of minimal amounts (1nM) of retinoic acid (SL^RA^) increased this fraction to more than 30%, which required activation of LIF and FGF but not of BMP signaling. Transdifferentiation of *Hdac3*^**-/-**^ cells is therefore driven by the same pathways that control progression of ICM into PrE *in vivo* (Hermitte and Chazaud, 2014; Morgani and Brickman, 2015).

To explore the ability of *Hdac3* mutants to mimic additional aspects of ICM development, we turned to a 3D culture system. After 3d in SL^RA^, single *Hdac3*^**-/-**^ but not *WT* cells formed spatially organized spheroids (**Figure 1G, S1L**): *Nanog* reporter-positive cells were enriched in the inside, while *Gata6* reporter-positive cells co-expressing Sox17 and showing polarized distribution of the apical PrE marker Dab2 (Hermitte and Chazaud, 2014) were on the outside. This resembles formation of the polarized epithelial PrE cell layer on the surface of the epiblast in E4.5 embryos and indicates spatial lineage segregation in *Hdac3*^**-/-**^ spheroids.

### Dax1 inhibits mESCs transdifferentiation into PrE similarly to Hdac3

To gain a more complete understanding of lineage restriction, we set out to determine the relationship of Hdac3 with two previously described inhibitors of PrE transdifferentiation, Dax1 and Prdm14 (Khalfallah et al., 2009; Ma et al., 2010; Zhang et al., 2014). Naïve and SL^RA^-exposed *Prdm14*^**-/-**^ cells generated in the G6C18 background were indistinguishable from *WT* controls. In contrast to *Prdm14*^**-/-**^ and similar to *Hdac3*^**-/-**^ cells, *Dax1* mutants showed increased *Nanog* reporter levels in 2iL, and approximately 30% of the cells expressed the *Gata6* reporter after 3d in SL^RA^ (**Figure 2A, S2A**-**C, Table S1**).

**Figure 2:**
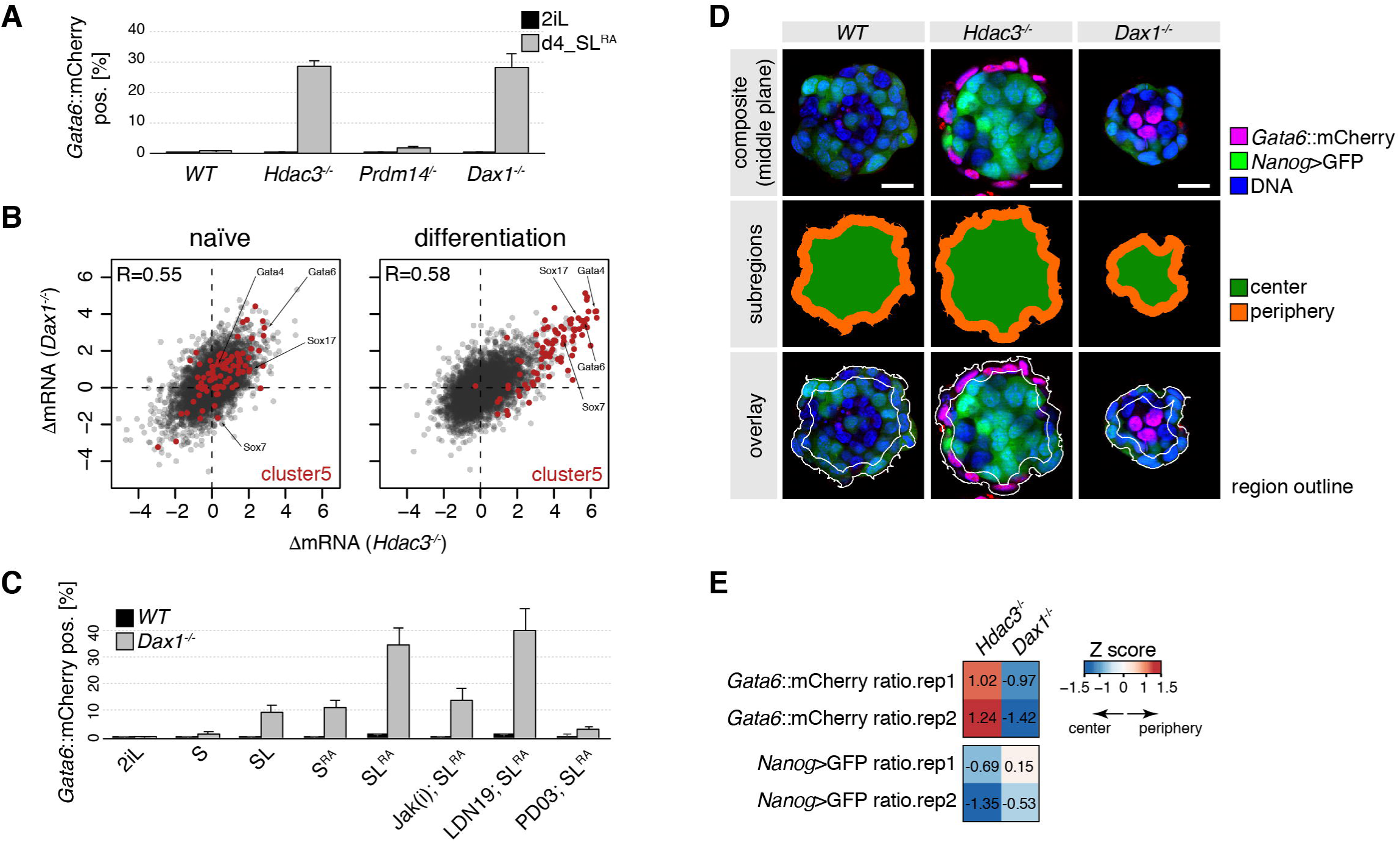
*Dax1* is required for lineage restriction and sorting. **(A,C)** Fraction of *Gata6*::mCherry positive cells in indicated genotypes and conditions. Average and SD of at least two independent clones. **(B)** Scatter plots of log2 fold change (FC) gene expression changes in *Hdac3*^**-/-**^ and *Dax1*^**-/-**^ mutants relative to *WT* cells in naïve mESCs (top) and after 2 d of differentiation (bottom); Cluster 5 genes are colored and selected PrE TFs labeled. **(D,E)** Quantification of spatial lineage segregation in spheroids. Representative spheroid segmentations of indicated genotypes (D). Z-score-normalized fluorescence distributions in mutants compared to *WT* spheroids (E). Positive Z-scores indicate peripheral, outside, and negative Z-scores central, inside, enrichment. Scalebar: 25μM.

RNAseq of *Dax1* mutants revealed that transcriptional changes in general correlated with those in *Hdac3* mutants in both, 2iL (R=0.55) and during differentiation (R=0.58) (**Figure 2B, Table S3**). Cluster 5 genes were partially deregulated in 2iLIF in both *Dax1*^**-/-**^ and *Hdac3*^**-/-**^ mESCs, indicating PrE-priming (**Figure 2B**). Also, the signaling pathway dependencies for PrE transdifferentiation were the same for *Dax1*^**-/-**^ and *Hdac3*^**-/-**^ cells (**Figure 1F, 2C, S2D**). In *Dax1* mutant 3D aggregates, however, peripheral enrichment of Gata6 reporter-positive PrE cells was perturbed when compared to *Hdac3* mutants (**Figure 2D,E, S2E, Table S3**), suggesting that *Dax1* has additional roles in spheroid self-organization.

We note that *Dax1* knockout and knockdown mESCs in SL have been described before, reporting apart from the induction of PrE markers (Fujii et al., 2015; Niakan et al., 2006; Zhang et al., 2014) lack of viability (Yu et al., 1998), loss of pluripotency (Khalfallah et al., 2009; Niakan et al., 2006), induction of 2-cell stage specific genes (Fujii et al., 2015), and multi-lineage differentiation (Khalfallah et al., 2009). We speculate that these discrepancies arise because absence of *Dax1* in metastable, but not naïve, conditions destabilizes pluripotency and results in or exacerbates culture heterogeneity.

### Hdac3 and Dax1 independently restrict PrE fate and antagonize Nr5a2 and Esrrb

PrE conversion upon loss of *Dax1* and *Hdac3* was qualitatively and quantitatively highly similar, suggesting that Dax1 and Hdac3 may act together, potentially in a protein complex. However, affinity-purification coupled to label-free quantitation (**Figure 3A, Table S3**) revealed that Hdac3 co-immunoprecipitated subunits of the Ncor1/Ncor2 complexes (Gps2, NCor1, NCor2, Tbl1x, Tbl1xr1), but not Dax1. *Vice versa*, Dax1 formed a complex with the nuclear receptors Nr5a2 and Esrrb, but not with Hdac3. To test if Hdac3 and Dax1 mechanisms of PrE repression were truly independent, we analyzed their genetic interaction by generating compound knock-out cell lines (**Figure S2B, Table S1**). After 3d in SL^RA^, PrE marker levels in *Dax1*^**-/-**^;*Hdac3*^***-*/-**^ double mutants were elevated two-to threefold compared to single mutants (**Figure 3B**). Notably, this additivity was due to a doubling of the fraction of cells expressing the *Gata6* reporter, reaching to more than 70% (**Figure 3C, S3A**). Dax1 and Hdac3 therefore act in parallel pathways that threshold the probability of single cells to exit pluripotency and activate the PrE program.

**Figure 3:**
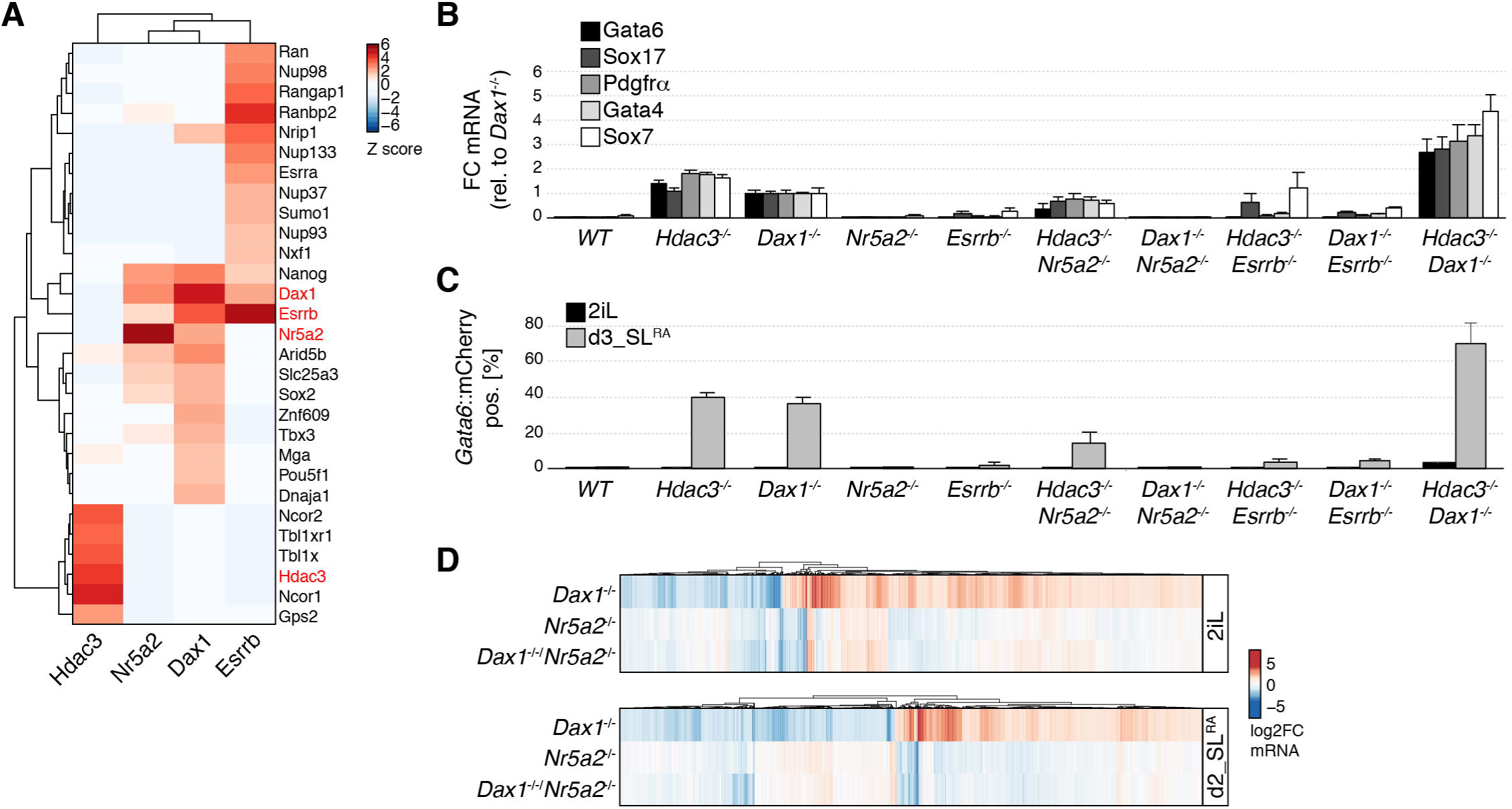
Hdac3 and Nr5a2/Dax1 are genetically and biochemically distinct pathways repressing PrE differentiation. **(A)** Z-scores of high-confidence interactors coIPed by Hdac3, Dax1, Nr5a2, and Esrrb (indicated in red) in naïve mESCs. **(B,C)** Expression of PrE markers relative to *Dax1*^**-/-**^ cells (B), and fraction of *Gata6*::mCherry positive cells after 3d in SL^RA^ (C) in indicated genotypes. Average and SD of three independent clones. **(D)** Gene expression changes in *Dax1*^**-/-**^, *Nr5a2*^**-/-**^ and *Dax1*^**-/-**^;*Nr5a2*^**-/-**^ cells compared to WT cells in 2iLIF (upper) and after 2d in SL^RA^ (lower).

Since Nr5a2 and Esrrb have been implicated in modulating PrE gene expression (McDonald et al., 2014; Uranishi et al., 2016), we decided to investigate their role in more detail. Consistent with the specific binding to Dax1, deletion of *Nr5a2* completely reverted upregulation of the *Nanog* and *Gata6* reporters and of PrE marker genes in *Dax1*^**-/-**^ cells, but only modestly in *Hdac3*^**-/-**^ cells (**Figure 3B,C, S2B, S3A,B, Table S1**). RNAseq in 2iL and SL^RA^ similarly showed that the majority of transcriptional changes induced by absence of *Dax1*, including those of cluster 5 genes, were reverted by co-deletion *of Nr5a2*, while loss of *Nr5a2* on its own caused comparatively minor defects (**Figure 3D, S3C,D, Table S3**). Therefore, Nr5a2 is the key interactor mediating Dax1 function. In contrast to *Nr5a2*, deletion of *Esrrb* partially suppressed the upregulation of the *Gata6* reporter and of PrE marker genes in both, *Dax1* and *Hdac3* mutants (**Figure 3B,C, S2B, S3A**), suggesting that *Esrrb* has non-specific functions downstream of *Dax1* and *Hdac3*. In summary, Dax1/Nr5a2 and Hdac3/Ncor1/Ncor2 form biochemically and genetically distinct complexes that antagonize PrE fate in mESCs.

### Gata6 enh^-45^ is a mechanistic target of Hdac3 and required for transdifferentiation

The transcriptional and phenotypic similarity of *Hdac3* and *Dax1* mutants indicates that they are independently acting components of the same regulatory network. To identify the mechanism of convergence, we exploited published Hdac3, Dax1 and Nr5a2 chromatin immunoprecipitation sequencing (ChIPseq) data (Atlasi et al., 2019; Beck et al., 2014; Żylicz et al., 2018), and determined cis-regulatory elements (CRE) activity changes by performing assays for transposase-accessible chromatin sequencing (ATACseq), and H3K27ac and H4K5ac ChIPseq in *Dax1*^**-/-**^ and *Hdac3*^**-/-**^ cells, respectively. Focusing on 29’969 regions bound by either Hdac3 or Dax1 (**Table S4**) we found that Dax1 occupancy was more similar to that of Nr5a2 (R=0.58) than of Hdac3 (R=0.11) (**Figure S4A**), which is consistent with distinct biochemical complexes. CRE activity changes in differentiating *Dax1* and *Hdac3* mutants, in contrast, correlated (**Figure S4B**), and clustering analysis (**Figure 4A**) identified two clusters of loci that were enriched for Hdac3 and Dax1 binding, and linked to CRE repression (cluster 2, decrease of accessibility, H3K27ac and H4K5ac) and activation (cluster 4, increase of accessibility, H3K27ac and H4K5ac). Notably, *Nr5a2* deletion reverted CRE activity changes in *Dax1* mutants (**Figure 4A**). Dax1/Nr5a2 and Hdac3 therefore bind to and regulate shared CREs.

**Figure 4:**
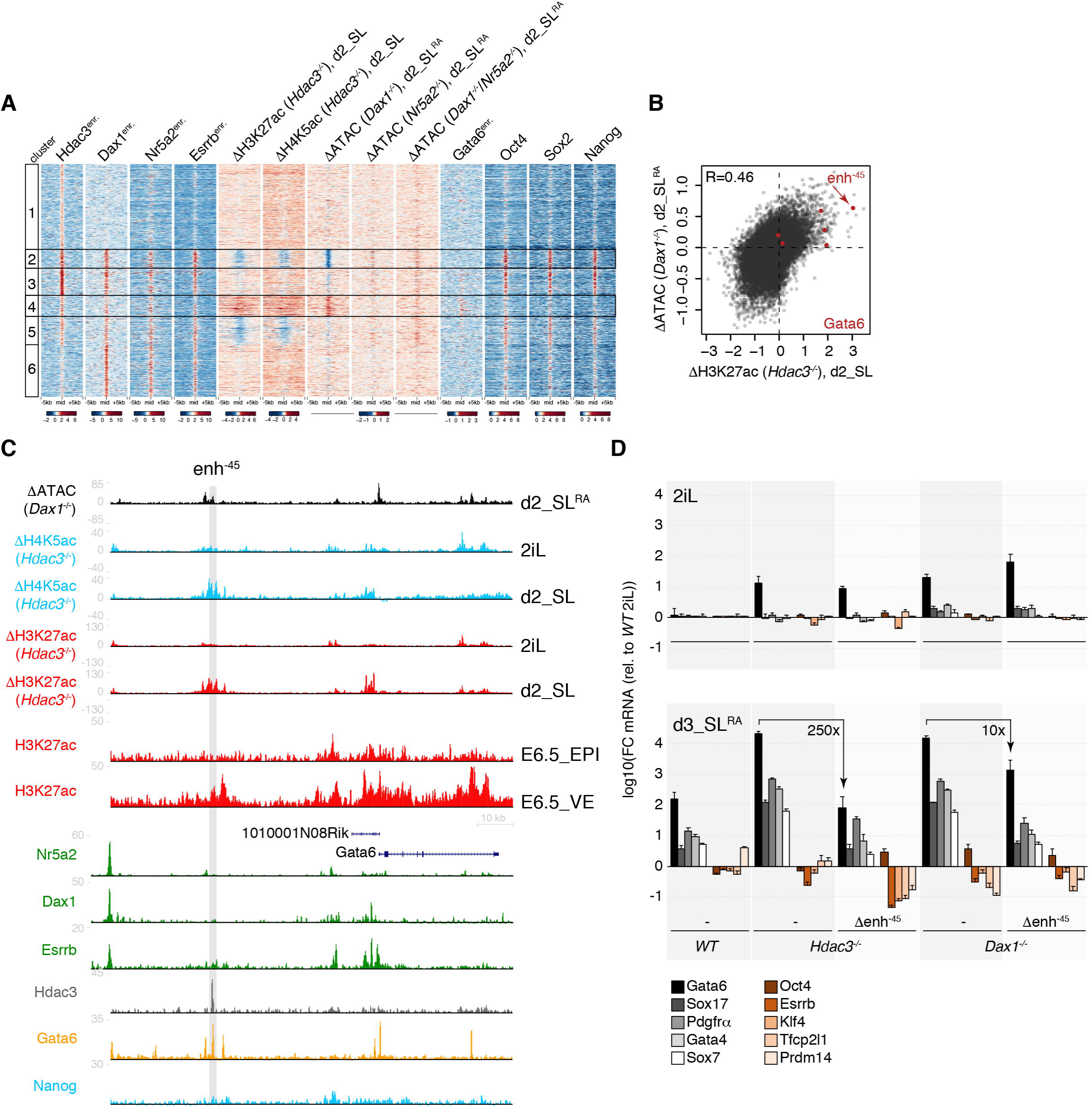
Hdac3 and Dax1 co-repress Gata6-bound CREs, including the essential Gata6 enh^-45^. **(A)** k-means clustered heatmap showing enrichment of TFs, and changes of chromatin marks or accessibility in indicated mutants relative to *WT* at Hdac3- and/or Dax1-bound regions. Repressed and induced clusters 2 and 4 are marked. **(B)** Scatterplot of log2 fold accessibility changes in d2_SL^RA^ *Dax1*^**-/-**^ cells and of log2 fold H3K27ac ChIP signal changes in d2_SL *Hdac3*^**-/-**^ cells relative to *WT* controls at Hdac3- and/or Dax1-bound regions. Regions colored in red are associated with Gata6. **(C)** Genome-browser view of the Gata6 locus showing accessibility and chromatin mark changes in indicated mutants, and of ChIPseq signals of indicated TFs and H3K27ac in E6.5 VE and EPI. **(D)** FC of PrE and naïve pluripotency markers relative to *WT* cells in 2iL of indicated genotypes in indicated conditions. Average and SD of at least three independent clones.

To determine the molecular features that are associated with co-regulated CREs, we scanned underlying TF motifs (**Figure S4C,D, Table S4**). This revealed enrichment of the GATA and OCT4-SOX2 consensus motifs at CREs that were activated and inactivated in mutant cells, respectively. Comparison with ChIPseq confirmed binding of Oct4, Sox2 and Nanog (Marson et al., 2008) to CREs that are ectopically silenced in naïve and differentiating *Hdac3* and *Dax1* mutants, and binding of Gata6 (Wamaitha et al., 2015) to CREs that are ectopically activated in differentiating but not naïve mutant cells data (**Figure 4A, S4E**). Hdac3 and Dax1 therefore converge on regulating CREs co-bound by core pluripotency TFs, and on repressing PrE-specific CREs that are occupied by Gata6 and activated during transdifferentiation.

As Gata6 overexpression is sufficient to convert mESCs into extraembryonic endoderm stem (XEN) cells *in vitro* (Shimosato et al., 2007), we hypothesized a causative role for Gata6 in the PrE transdifferentiation of *Dax1* and *Hdac3* mutants. This possibility is supported by specific upregulation of Gata6 in naïve single and compound mutants (**Figure S4F**) and by the fact that a region 45 kb upstream of the *Gata6* gene (enh^-45^) was amongst the most strongly activated CREs in mutant cells both during differentiation and in 2iL (**Figure 4B,C, S4G**), and decorated with H3K27ac in the E6.5 VE but not EPI (Xiang et al., 2019) (**Figure 4C**). enh^-45^ is bound by Hdac3 and Gata6, but not Dax1, raising the possibility that it is a relevant direct Hdac3 target. To test this, we deleted the 500bp region that is occupied by Hdac3 (Δenh^-45^) (**Table S1**), which fully suppressed activation of the *Gata6* reporter and reduced PrE marker expression in Δenh^-45^;*Hdac3*^**-/-**^ and Δenh^-45^;*Dax1*^**-/-**^ cells to levels observed in the respective single mutants upon SL^RA^ transition (**Figure 4D, S4H,I**). Reversion of *Gata6* transcription was more efficient in *Hdac3* than *Dax1* compound Δenh^-45^ mutants, but absent in naïve mESCs (**Figure 4D**). Surprisingly, removal of enh^-45^ triggered downregulation of the *Nanog* reporter and of the naïve pluripotency markers Klf4, Esrrb, Tfcp2l1 and Prdm14 in *Hdac3* but not *Dax1* mutants in SL (**Figure 4D, S4H**). This indicates additional roles of Hdac3 in the stability of the pluripotent GRN that are masked by enh^-45^ regulation and not shared with Dax1. We conclude that direct and indirect suppression of enh^-45^ by Hdac3 and Dax1, respectively, is crucial for repressing Gata6 transcription and, consequentially, PrE transdifferentiation upon SL transition.

## DISCUSSION

Here we show that the transcriptional repressors Hdac3 and Dax1 set the lineage barrier that prevents mESCs from transdifferentiating into PrE upon exposure to developmental signals. In naïve conditions, *Hdac3* and *Dax1* mutant mESCs are stable, allowing us to exploit the transition from 2iL to SL conditions as paradigm for PrE conversion (Schröter et al., 2015). Adaption to 3D results in formation of *Hdac3* mutant spheroids in which pluripotent and PrE lineages are spatially segregated. In *Dax1* mutants, segregation is perturbed. This setup is therefore amenable to dissecting the molecular requirement underlying PrE differentiation and sorting.

Hdac3 and Dax1 cooperatively inhibit PrE transdifferentiation. Notably, conversion of naïve *Hdac3*^**-/-**^ and *Dax1*^**-/-**^ mESCs occurs without overexpression of PrE-specifying TFs, such as Sox17 (McDonald et al., 2014), Gata4 or Gata6 (Fujikura et al., 2002), or use of selective culture conditions that confer expanded potential (Sozen et al., 2019; Yang et al., 2017) or select the survival and/or proliferation of PrE cell types (Anderson et al., 2017), but upon transition to SL, which supports both, EPI and PrE fates. The pluripotent gene regulatory network is therefore permissive to lineage switching in the absence of ectopic reprogramming TFs or developmental reversion.

Dax1 and Hdac3 act in independent protein complexes that converge on repressing *Gata6*. We show that lineage conversion in the absence of *Dax1* and *Hdac3* requires activation of a single *Gata6* enhancer, enh^-45^, that is directly targeted by Hdac3. Recruitment of epigenetic regulators, such as Sin3a (Mall et al., 2017) or Groucho (Kutejova et al., 2016; Muhr et al., 2001) is associated with the repression of alternate lineages, but this study – to our knowledge for the first time – links the physical binding and enzymatic activity of a transcriptional repressor, Hdac3, at a single CRE with suppression of transdifferentiation. enh^-45^ is occupied by Gata6 during transdifferentiation, indicating a positive feedforward loop that is characteristic of cell type-specifying TFs (Crews and Pearson, 2009) and can stabilize cell identity (Leyva-Díaz and Hobert, 2019). Our findings therefore suggest that the continuous silencing of autoregulated enhancers of competing master TFs is an endogenous mechanism for maintaining lineage restriction.

We find that Hdac3 and Dax1 repress transdifferentiation by modulating the pluripotent GRN network, specifically Nr5a2 and Esrrb, which are TFs that regulate mESC self-renewal (Festuccia et al., 2020) and iPSC reprogramming (Feng et al., 2009; Guo and Smith, 2010). Nr5a2 is a well-established interactor of Dax1 (Sablin et al., 2008). Co-deletion of *Nr5a2* rescues PrE conversion and most transcriptional and epigenetic alterations in *Dax1* but not *Hdac3* mutants. Therefore, Nr5a2 and Dax1 operate as a functional unit where Nr5a2 is the transcriptional activator and Dax1 the repressor. The direct targets of Nr5a2 driving PrE differentiation remain unclear, since Nr5a2 and Dax1 do not bind to enh^-45^. Lack of *Esrrb*, in contrast, partially blocks PrE transdifferentiation of *Dax1* and *Hdac3* mutants. Although Esrrb interacts and shares genomic binding with Dax1 and Nr5a2 (**Figure 3A, 4A, S4A**), incomplete phenotypic rescue and lack of specificity for *Dax1* suggests a role of Esrrb in PrE conversion beyond the Nr5a2/Dax1 axis, e.g. by binding (**Figure 4C**) and activating the Gata6 promoter (Uranishi et al., 2016) or, indirectly, by stabilizing pluripotency (Martello et al., 2012). Taken together, we propose that Nr5a2 and Esrrb support both the EPI and the PrE fate within a coherent TF network that integrates extrinsic signals with transcriptional repressors to maintain lineage choice.

In summary, we uncover how the pluripotent lineage is safeguarded in mESCs. We find that both, Gata6 and Nanog, are induced in naïve *Hdac3*^**-/-**^ and *Dax1*^**-/-**^ mESCs, and that enh^-45^ is not occupied by Nanog (**Figure 4C**). This is inconsistent with mutual inhibition of Gata6 by Nanog and indicates that maintenance of the EPI-PrE segregation is independent of direct TF antagonism. We, instead, suggest operation of a GRN that (a) epigenetically silences Gata6, the master TF of the competing PrE fate, and (b) balances the activities of lineage-divergent TFs, such as Nr5a2 and Esrrb (**Figure S4J**). We speculate that this regulatory network is established during developmental progression of pluripotency to stabilize the initially labile lineage segregation set by antagonism between Nanog and Gata6.

## ACKNOWLEDGMENTS

We thank H.Kohler (FMI) for cell sorting, Karoline Guja for technical support, V. Iesmantavicius (FMI) for help with computational analysis of mass spectrometry data, Min Jia and Jeff Chao (FMI) for providing LIF, and Austin Smith (Wellcome-MRC Cambridge Stem Cell Institute) for providing TNG-A cells. We are grateful to F. Mohn, A.H.F.M. Peters, D. Schuebeler (FMI) and S. Stricker (University Munich) for comments on the manuscript. D.O. was supported by EMBO (ALTF 1632-2014) and Marie Curie Actions (LTFCOFUND2013, GA-2013-609409). Research in the lab of J.B. is supported by the Novartis Research Foundation.

## AUTHOR CONTRIBUTIONS

D.O. conceived the study and performed experiments, P.P. and J.B. bioinformatical analysis, M.R. RT-qPCRs, D.H. mass spectrometry, and I.L. spheroid image analysis. S.S. supervised sequencing library generation and sequencing, and M.B.S. bioinformatical analysis. D.O. and J.B. wrote the paper.

## MATERIALS AND METHODS

### Mouse ESCs

Male TNG-A mESCs where a GFP-IRES-Puromycin-N-acetyltransferase is knocked into one of the Nanog alleles (Chambers et al., 2007) are a gift from Austin Smith (Wellcome – MRC Cambridge Stem Cell Institute and Living Systems Institute, University of Exeter).

### Cell culture

mESCs were cultured on gelatin-coated plates in N2B27 medium (DMEM/F12 medium (Life Technologies) and Neurobasal medium (Gibco) 1:1, N2 supplement 1/200 (homemade), B-27 Serum-Free Supplement 1/100 (Gibco), 2mM L-glutamine (Gibco), 0.1mM 2-mercaptoethanol (Sigma)), 1µM PD0325901, 3µM CHIR99021 (Steward lab, Dresden), and mLIF (Smith lab, Cambridge; Chao lab, FMI). For medium switch and differentiation experiments cells were washed with PBS, detached in Accutase (Sigma), centrifuged at 300g for 3 minutes (min) in DMEM/F12 with 0.1% BSA (Gibco), resuspended in the appropriate medium, counted using Fast Read 102 counting chambers (Kova), diluted and plated. Cells were plated in Serum LIF (SL) medium (GMEM (Sigma), 10% fetal bovine serum (Sigma), 1mM sodium pyruvate (Gibco), 2mM L-glutamine (Gibco), 0.1mM non-essential amino acids (Gibco), 0.1mM 2-mercaptoethanol (Sigma), and mLIF), plus Retinoic acid (1nM, Sigma), JAK Inhibitor I (1µM, Calbiochem) and LDN193189 (100nM, Sigma), PD0325901, where specified, at 10’000 cells/cm^2^ on gelatin-coated plates, or in EpiLC medium (N2B27 medium, 20 ng/ml activin A and 12 ng/ml bFGF (Smith lab, Cambridge), and 1% KSR (Life Technologies)) at 25’000 cells/cm2 on fibronectin-coated plates. To generate spheroids, 1200 cells were plated as described above in 1.5 mL SL plus 1nM Retinoic acid in a well of an AggreWell 400 plate (Stemcell Technologies), prepared with anti-adherence rinsing solution (Stemcell Technologies) according to manufacturer’s instructions, and incubated for 3 days.

TNG-A mESCs stably transfected with pPB-LR51-EF1a-bsdr2ACas9 (modified from (Koike-Yusa et al., 2013)) were reverse transfected with U6>sgRNA plasmids (George Church, Addgene plasmid #41824) as follows: 3µL of Lipofectamin2000 (Life Technologies) were mixed with 250µL of OPTIMEM (Gibco) and incubated for 5 min at RT, 350ng of sgRNA plasmid diluted in 250µL of OPTIMEM were added to the Lipofectamin2000 mix and incubated for 30 min at room temperature (RT), the transfection mix was added to freshly resuspended 200’000 cells in 2mL of medium in a well of a 6-well-plate. The next day medium was changed. 3 days after transfection, single cells were sorted in 96-well plates for clonal isolation. Parental cell line, sgRNA sequences, genotyping strategy, and sequencing results/western blots for every knock-out clone used in this study are detailed in **Table S1** and supplementary figure panels. The Hdac3Y118F knock-in cell lines were generated by reverse transfection as above of 400ng Hdac3_gRNA7 plasmid and 10pmol of oligo Hdac3YF_ki_f (for clone 1034-15) or oligo Hdac3YF_ki_r (for clone 1035-6). The Gata6::mCherry knock-in cell line G6C18 was generated by reverse transfection as above of 300ng Gata6-C gRNA#1 plasmid and 800ng of Gata6_3xGS_mCherry homologous recombination template. sgRNAs, oligonucleotide sequences, genotyping strategies, and sequencing results for knock-in clones are detailed in **Table S1**.

### siRNA transfections

Cells were reverse transfected in 2iLIF as follows: 2.4µL of LipofectamineRNAiMAX (Life Technologies) were mixed with 200µL of OPTIMEM (Gibco) and incubated for 5 min at RT, 2.4µL of 20µM Ncor1 siRNA mix and Ncor2 siRNA mix or AllStars Neg. Control siRNA (FlexiTube GeneSolution, Qiagen) diluted in 200µL of OPTIMEM were added to the LipofectamineRNAiMAX mix and incubated for 30 min at RT, the transfection mix was added to freshly resuspended 100’000 cells in 1.6mL of medium in a well of a 12-well-plate. The next day the cells were detached and transferred to fresh 2iLIF or Serum LIF. siRNA sequences are displayed in **Table S1**.

### Western blotting

Cells were lysed in 1x RIPA buffer (50mM Tris pH7.5, 150mM NaCl, 1% IGEPAL, 0.5% Na Deoxycholate, 0.1% SDS, 2mM EDTA), protein concentration was determined by the BCA protein assay (Pierce), and 10µg of protein per sample were loaded on 10% polyacrylamide gels. Primary antibodies used: anti-HDAC3 antibody, ab7030, by abcam, 1/5000; DAX-1/NR0B1 antibody (mAb), Clone: 1DA-2F4, Active Motif, Catalog No:39983, 1/10’000; Human ERR beta/NR3B2 Antibody, PP-H6705-0, by R&D Systems, 1/10’000.

### Quantitative PCR (qPCR)

RNA was purified with the RNeasy Mini Kit with on-column DNase digest (Qiagen). 1µg of RNA was reverse transcribed using SuperScript III Reverse Transcriptase (LifeTechnologies) and qPCR was performed with the TaqMan Fast Universal PCR Master Mix (Thermo Fisher) and the following TaqMan probes: GAPDH (4352339E), Gata6 (Mm00802636_m1), Gata4 (Mm00484689_m1), Pdgfra (Mm00440701_m1), Sox7 (Mm00776876_m1), Sox17 (Mm00488363_m1), Oct4 (Mm03053917_g1), Esrrb (Mm00442411_m1), Klf4 (Mm00516104_m1), Tfcp2l1 (Mm00470119_m1), Prdm14 (Mm01237814_m1).

### Co-immunoprecipitations (IPs)

Endogeneous Hdac3 was IP’ed from nuclear extract of G6C18 cells in 2iLIF using anti-HDAC3 antibody (ab7030, by abcam) and rabbit normal IgGs (Santa Cruz) as control. N-3xFLAG-3GS-Dax1, Nr5a2-3GS-3xFLAG-C, and Esrrb-3GS-3xFLAG-C cloned in pPB-CAG-DEST-pgk-hph (Betschinger et al., 2013) were expressed in G6C18 cells in 2iLIF and IP’ed from nuclear extracts using anti-FLAG M2 antibody (Sigma) using un-transfected cells as control. Briefly, cells from a confluent T-75 flask were harvested and nuclei isolated by hypotonic buffer (10mM TrisHCl pH7.5, 10mM KCl, 1mM dithiothreitol (DTT), 0.5% IGEPAL) for 20 min on ice. Nuclei were lysed by rotation for 90 min at 4°C in 1mL of lysis buffer (20mM TrisHCl pH7.5, 100mM KCl, 1.5mM MgCl2, 1mM DTT, 10% glycerol, 1x Protease Inhibitor Tablet (Roche), 2x Phosphatase Inhibitor Tablets (Roche), 0.5% Triton X-100, 250 units/mL Benzonase (Sigma)). Clarified lysates were incubated with 1.2µg of antibody pre-bound to 5µL protein Dynabeads (Life Technologies) for 6 hours (h) at 4°C. Beads were washed 4 times with wash buffer (20mM TrisHCl pH 7.5, 150mM NaCl, 0.5% IGEPAL) and 3 times with wash buffer without detergent, and digested with 0.2µg Lys-C (WAKO) and 0.2µg modified porcine trypsin (Promega) before subjecting to mass spectrometry as described before (Villegas et al., 2019).

### Flow cytometry

Cells were washed with PBS, detached in Accutase, centrifuged at 300g for 3 min in DMEM/F12 with 0.1% BSA, resuspended in DMEM/F12 with 0.1% BSA, run on an LSRII SORP Analyzer (Becton Dickinson), and analyzed using FlowJo (FlowJo, LLC). Quantification for Gata6::mCherry positive cells (**Figure 1F, 2A, 2B, 3C, S4I**) were performed on experimental triplicates, with gates shown in fluorescence intensity plots. Nanog>GFP intensities (**Figure S2C, S3B**) are the average of geometric GFP mean intensities of experimental triplicates.

### High-throughput imaging of spheroids

To fix the spheroids, 1mL of medium was removed from each well and 500µL of 8% PFA (Alfa Aesar) in PBS were added to the remaining 500µL of medium and incubated for 20 min at 37°C. The spheroids were then harvested with a 1000µL pipette, washed twice with PBST (PBS-0.1% Tween-20 (Sigma)) and once with PBST-Hoechst 33342 (1/5000, Life Technologies), transferred in PBS to a µCLEAR 96-well plate (Greiner Bio-One), and imaged on a Yokogawa CV7000s high throughput confocal microscope at 40x magnification. Images were acquired in confocal mode as z-stack multiplane images over a z distance of 100 µm with 5 µm step width. Subsequently, images were stitched to generate a single image per z-plane, channel and well, which was used for object segmentation and feature extraction.

### Antibody staining of spheroids

Fixed spheroids were incubated in blocking solution (3% donkey serum (Sigma), 1% BSA in PBST) for 1 h at RT, stained overnight at 4°C with antibodies (anti-Dab2 monoclonal antibody (D709T), #12906S Cell Signaling Technology, 1/200; Human SOX17 Antibody, #AF1924 R&D Systems, 1/200) in blocking solution, washed 3 times with PBST, stained with secondary antibodies (goat anti-rabbit Alexa Fluor 647 and donkey anti-goat Alexa Fluor 647, 1/500, ThermoFisher) in PBST for 1h at RT, washed twice with PBST and once with PBST-Hoechst 33342, transferred in PBS to a µCLEAR 96-well plate (Greiner bio-one), and imaged on a Zeiss LSM710.

### Native histone ChIPseq

Nuclear lysates were prepared from 10 million cells as for coIPs above. Clarified lysates were incubated with 4µg of primary antibody for 2h and then SDS concentration was brought to 0.1% and EDTA to 5mM to inactivate benzonase. Lysates were clarified again and 40µL of protein G Dynabeads were added and rotated for 6h at 4°C. Beads were washed twice with RIPA buffer, four times with RIPA buffer with 500mM NaCl, and once with 10mM Tris-HCl pH8.0 with 1mM EDTA. DNA was eluted by proteinase K (Macherey-Nagel) treatment in 0.5% SDS for 3h at 55°C and purified with Qiagen MinElute PCR Purification Kit. Primary antibodies used: H4K5ac (Merck-Millipore, 07-327), H3K27ac (Active Motif, 39135). Sequencing libraries for biological duplicates of each assayed histone modification and input per condition were prepared using the ChIP-Seq NEB Ultra (Dual Indexes) Kit (New England Biolabs) and sequenced on an Illumina HiSeq2500 (50 bp single-end reads). In total ∼1.05 billion reads (accounting to ca 45 million reads per replicate) were generated. After demultiplexing, reads were aligned against the mouse genome (GRCb38/mm10) using the QuasR Bioconductor library (Gaidatzis et al., 2014) with default parameters. The overall alignment rate was ∼95%.

### ATACseq

ATAC-seq was performed as describe before (Buenrostro et al., 2015). Briefly, freshly isolated nuclei from 50’000 cells were subjected to transposition using the Nextera Tn5 Transposase for 30’ at 37°C. DNA was purified with the Qiagen MinElute PCR Purification Kit and amplified for a total of 12 cycles with the Q5 PCR mix (NEB). Libraries of experimental triplicates were purified with AMPure-XP beads (Beckman Coulter) and sequenced on an Illumina NextSeq500 machine (75 bp paired-end reads).In total ∼0.96 billion reads (accounting to ca 80 million reads or 40 million read-pairs per replicate) were generated. After demultiplexing, reads were aligned against the mouse genome (GRCb38/mm10) using the QuasR Bioconductor library (Gaidatzis et al., 2014) with default parameters. The overall alignment rate was ∼94%.

### RNAseq

RNAseq of biological triplicates was performed using x cells. RNA was purified with the RNeasy Mini Kit with on-column DNase digest (Qiagen), RNAse treatment etc. Libraries were prepared using TruSeq mRNA Library preparation kit (Illumina) and sequenced on an Illumina HiSeq2500 machine (50 bp single-end reads). In total ∼1.6 billion reads (accounting for ca 30 million reads per replicate) were generated. After demultiplexing, reads were aligned against the mouse genome and guided by transcriptome annotation (GRCb38/mm10, GENCODE release M4) using STAR (Dobin et al., 2012) version 2.5.0 with command line parameters:

*--outSJfilterReads Unique --outFilterType BySJout --outFilterMultimapNmax 10 -- alignSJoverhangMin 6 --alignSJDBoverhangMin 3 –outFilterMismatchNoverLmax 0*.*1 --alignIntronMin 20 --alignIntronMax 1000000 --outFilterIntronMotifs RemoveNoncanonicalUnannotated --seedSearchStartLmax 50 --twopassMode Basic*

The overall alignment rate was consistently ∼90%.

## BIOINFORMATICS

### Integration of external embryonic RNAseq datasets and projection on common subspaces

**Integration of external embryonic RNAseq datasets and projection on common** 4 **subspaces**
| accession | RNAseq description | reference |
| --- | --- | --- |
| E-MTAB-2958 | mouse embryonic development | (Boroviak et al., 2015) |
| GSE100597 | mouse embryonic development | (Mohammed et al., 2017) |
| GSE77783 | endoderm from naïve pluripotency | (Anderson et al., 2017) |

GRCm38 alignments from (Boroviak et al., 2015) were downloaded (E-MTAB-2958) and exonic gene counts were obtained on the GENCODE M4 transcriptome annotation using the GenomicFeatures Bioconductor package (Lawrence et al., 2013). Single-cell RNAseq filtered count matrices from (Mohammed et al., 2017) were downloaded (GSE100597) and gene annotation was lifted from ENSEMBL to GENCODE M4 discarding ambiguous mappings. Assignments of individual cells to specific lineages were obtained through personal communication with the study authors and bulk gene counts for each stage and lineage were calculated by aggregating the corresponding single-cell data. Gene counts from all datasets were library normalized and converted to CPMs. Microarray data from (Anderson et al., 2017) were downloaded (GSE77783) and analyzed using the limma Bioconductor package (Ritchie et al., 2015). Pairwise correlation values (**Figure 1B, S1B, S1D**) between individual samples of two datasets were calculated after gene-wise mean normalization of the log2 gene expression values.

For the common subspace projection including datasets from different labs (**Figure 1C, S1C**), we converted gene expression values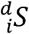of a sample *i* in dataset *d* to log2 fold gene expression changes (*S*’) relative to a matching reference sample *R* (*WT* 2iL and ESC (Boroviak et al., 2015), and *WT* d2_EpiLC and E6.5_EPI (Mohammed et al., 2017)):

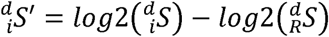

We used the stepwise estimation of common principal components (Trendafilov, 2010), implemented in the CRAN cpca package to obtain a common subspace projection for each pairwise comparison. cpca takes as input the sample covariance matrices of each dataset and performs a joint dimension reducing transformation under the assumption that the eigenvector space is identical across datasets. In each pairwise comparison only the intersection of the approximately top 5000 most variable genes, according to a within-dataset mean-variance trend fit for RNAseq experiments and a log2FC-based selection for microarray experiments, was used (provided in **Table S2**) for sample covariance matrix estimation.

### Differential gene expression analysis and gene clustering

Differential gene expression (**Figure 2B, 3D, S3C**) was calculated using the Bioconductor edgeR package (McCarthy et al., 2012; Robinson et al., 2009) (provided in **Table S3**). Briefly, for each experimental design a common negative binomial dispersion parameter was estimated using the *estimateGLMCommonDisp* function and a negative binomial generalized log-linear model was fit per gene using the *glmFit* function with a prior count of 1 in order to shrink log-fold change (LFC) effect sizes of lowly expressed genes towards zero. Reported p-values are the Benjamini & Hochberg adjusted estimates for multiple comparisons.

Clusters of genes with consistent expression profiles (**Figure 1D**) were identified with k-means clustering using as input features the gene expression log2FC of *WT* and *Hdac3*^**-/-**^ TNG-A derived clones in d2_SL and d2_EpiLC relative to *WT* TNG-A 2iL cells, focusing on the approximately top 5000 most variable genes as described above, and applying a pseudo-count of 1 to shrink effect sizes of lowly expressed genes. Only genes with a maximum absolute log2FC >1.5 in any of the four contrasts were considered (**provided in Table S2**). The calculation was carried out using the base R *kmeans* function implementation with *centers*=8 and 200 random starts. Analyses of enriched gene sets (**Figure S1F**) was performed using DAVID (Huang et al., 2008) for GO terms of biological processes.

### Peak calling

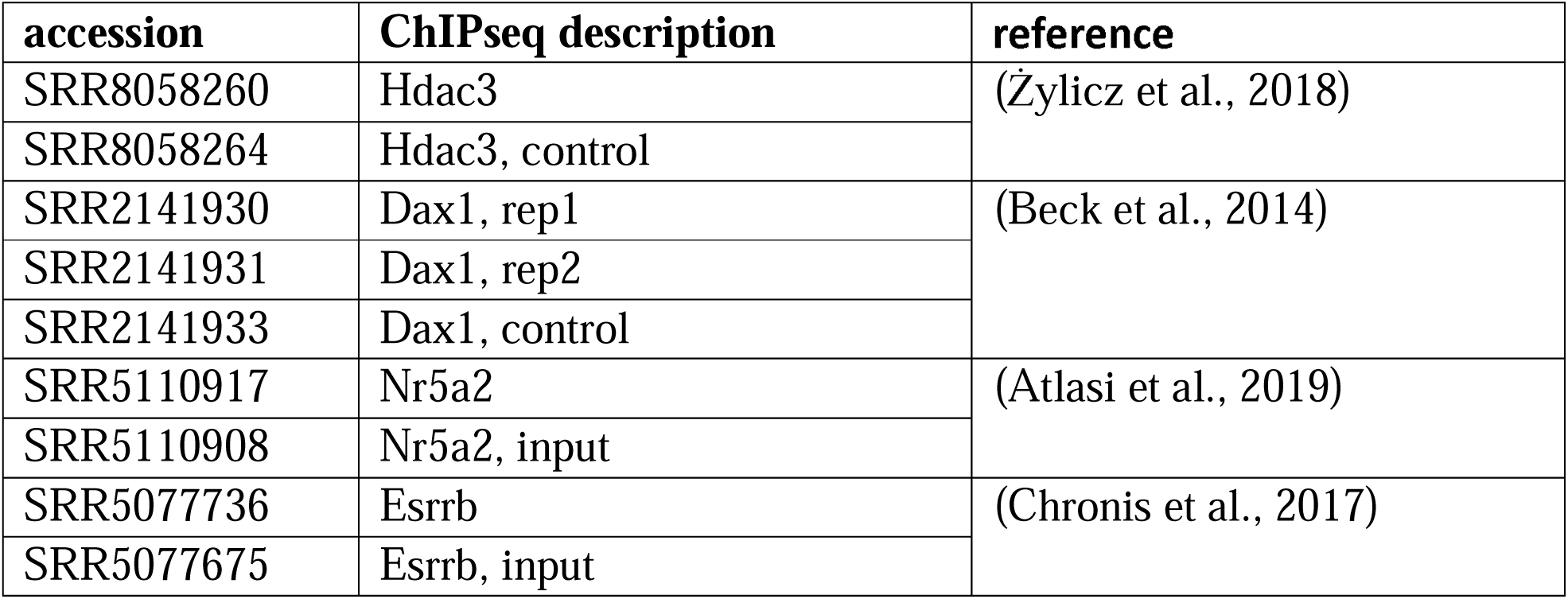
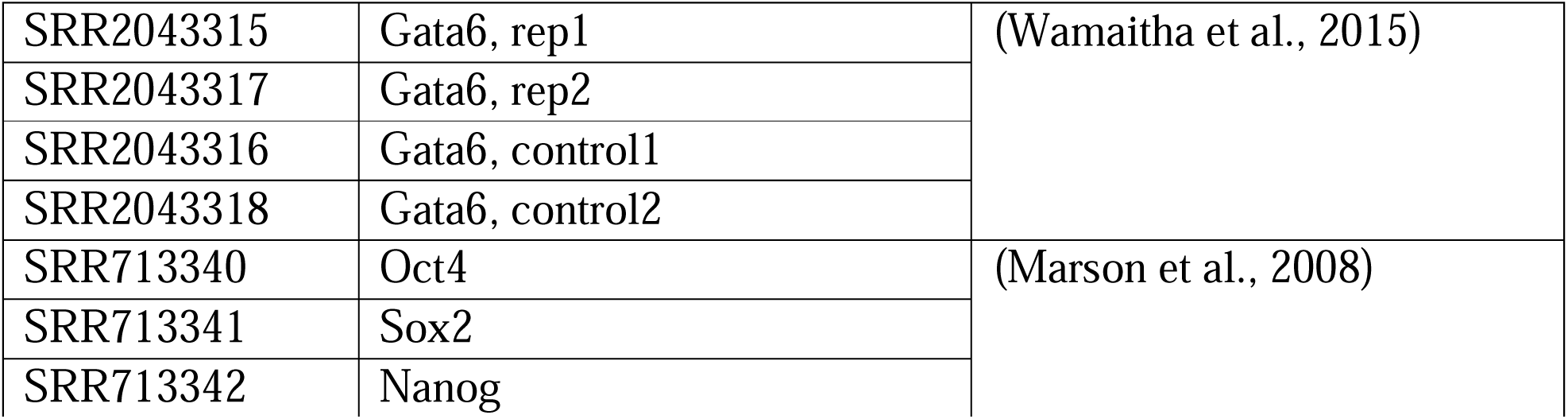

ChIP-seq data were realigned to GRCb38/mm10 using the QuasR Bioconductor library (Gaidatzis et al., 2014) with default parameters. Dax1 and Hdac3 peaks of significantly high read density, taken as a proxy of protein binding, were called using the csaw software (Lun and Smyth, 2015a) and according to the workflow outlined in (Lun and Smyth, 2015b). Briefly, the average fragment length was computed (function *correlateReads*) and reads were counted in windows of width w=78bp both for the precipitates and the input (where available). Windows of significant enrichment were filtered against a local background in regions of 5Kbp centered around the 78bp windows (3-fold enrichment, function *filterWindowsLocal*), and against the matching control sample (3-fold enrichment, function *filterWindowsControl*). Finally, nearby identified peaks, closer than 750nt apart, were merged (function *mergeWindows*). Blacklisted regions that often yield artifactual high signal in ChIP-seq experiments were obtained from the ENCODE project (https://www.encodeproject.org/files/ENCFF547MET/) and excluded from all read-counting operations (read parameter *discard*). Peaks were assigned to their proximal-most genes and distances to transcription start-sites (TSSs) were calculated according to the GENCODE release M4 annotation. A peak was considered as a “promoter peak” if it was within 1kbp distance of an annotated TSS.

### Differential ChIPseq/ATACseq enrichment

ChIPseq and ATACseq signal was quantified in 1kb regions at the 29969 peaks that are the union of Hdac3 (24249) and Dax1 (10671) peaks (**provided in Table S4**). In all cases, differential ChIP-seq or ATACseq enrichments (**Figure 4A, 4B, S4A-C, S4E, S4G**) between condition are stated, the values refer to changes in library normalized read counts, averaged, when available, over biological replicates. enr: denotes enrichment over control or input.

Clusters of peaks (**Figure 4A**) were identified with k-means clustering using as input features the enrichment of Hdac3 and Dax1 ChIPseq over controls, respectively, and differential H3K27ac ChIPseq and ATACseq signals in *Hdac3* and *Dax1* mutants, respectively, compared to *WT* cells after 2d of differentiation in SL. The calculation was carried out using the base R *kmeans* function implementation with *centers*=6 and 200 random starts. The Bioconductor ComplexHeatmap package (Gu et al., 2016) was used to plot enrichments in 10kb, peak-centered windows.

**Figure 4C** was assembled using the UCSC genome browser interface (https://genome.ucsc.edu).

### Motif enrichment analysis

Motif enrichment (**Figure S4D**) was performed as described before (Barisic et al., 2019). Briefly, the 29969 Hdac3/Dax1 peak union set was grouped based on differential H3K27ac and ATAC signal in mutants compared to *WT* cells after 2d of differentiation in SL into 9 bins with 1319 peaks each (**Figure S4C**). Peaks with absolute log2FCs less than 0.5 were grouped into a single bin (unchanged CRE activity). Positional weight matrices (PWMs) for 519 vertebrate transcription factors available from the JASPAR database through the Bioconductor JASPAR2016 package (Mathelier et al., 2015) were used to determine motif enrichment in each bin with the findMontifsGenome.pl script of the HOMER package (Heinz et al., 2010) using all other bins as background and parameters *-nomotif -size 500*. A cutoff for the expression of TFs in ESCs and during differentiation, and for the enrichment significance of their motifs was employed. Reported enrichment significance values are FDRs corrected for multiple testing.

### Mass spectrometry

Relative protein quantifications of three independent affinity-purifications (1: experimental duplicates of anti-Dax1, -Esrrb and -Nr5a2 coIPs, 2: experimental triplicates of anti-Dax1, -Esrrb, -Nr5a2 and -Hdac3 coIPs, 3: experimental triplicates of anti-Hdac3 coIPs) were merged and missing values filled in with a 1.8 fold downshift and a 0.3 fold distribution width of the actual distribution of detected proteins, as described (Tyanova et al., 2016). Fold-change enrichments were calculated over the respective experimental negative controls, and Z-scores per each coIP and experiment determined (provided in **Table S3**). Only proteins with Z-scores>2 in both independent affinity-purifications are considered (**Figure 3A**).

### Segmentation and quantification

Object segmentation was performed using image processing libraries available for Python 3. To identify the middle plane of spheroids segmentation masks were first created for the maximum intensity projections (MIPs) of the DNA channel by automated thresholding with 2-class Otsu algorithm. Entire z-stacks were then cropped to segmentation outlines obtained from the MIP segmentation. Subsequently, sum intensity of DNA staining in each plane was measured and used to identify the z-plane with maximum sum intensity of DNA signal, corresponding to the middle plane of the object. Segmentation masks were then adjusted in-plane by automated thresholding and used for subregion analysis. Each segmentation mask of the object middle plane was subdivided in 2 regions: A 50 pixel-wide concentric region along the object periphery and a central region, ranging from the object center to the border of the peripheral region. To exclude debris, objects where the central subregion was less than 30% of the object area were eliminated from the analysis. Areas and mean intensities of the Nanog>GFP and Gata6::mCherry channels were then separately calculated for the two regions, and used for quantitative analysis. Distribution of intensity between the peripheral and central regions was calculated as the ratio of mean intensity in the peripheral and central region (**Figure S2E**, provided in **Table S3**). Extracted features describing object area and intensity were normalized to the WT controls within corresponding biological replicates using z-score transformation and unified into a cross-comparable dataset (**Figure 2E**, provided in **Table S3**).

### Quantification and Statistical analysis

Details for quantification and statistical analysis are specified in the figure legends, including number of biological replicates. Data is presented as the average and standard deviation.

## DATA AND SOFTWARE AVAILABILITY

All sequencing data have been deposited at ArrayExpress (E-MTAB-9446, E-MTAB-944, E-MTAB-0450 and E-MTAB-9453).

## SUPPLEMENTAL FILES

**Figure S1:**
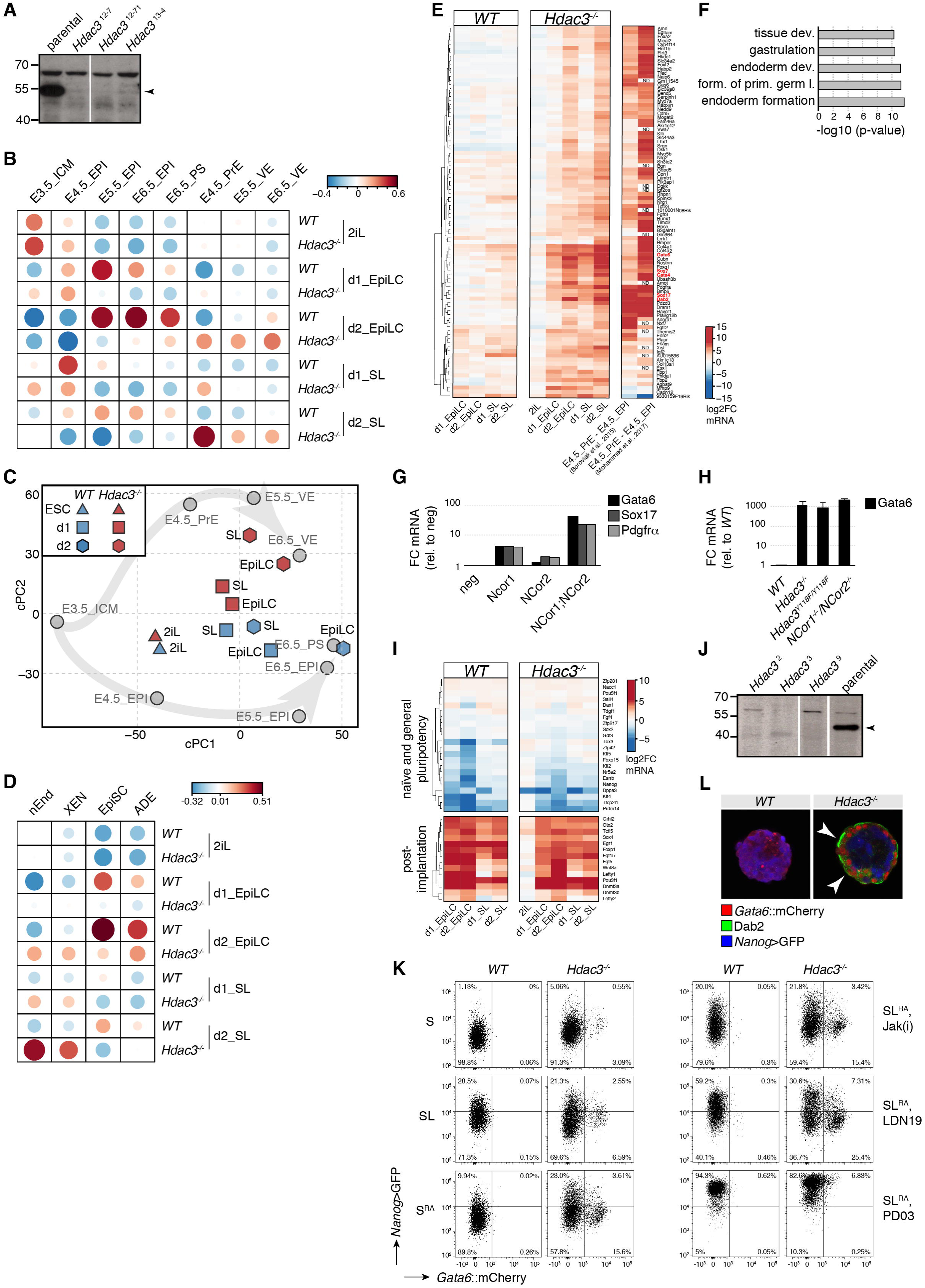
**Transcriptomics of *Hdac3*^-/-^ cells. Related to Figure 1.** **(A)** Anti-Hdac3 western-blot of naïve Hdac3^**-/-**^ TNG-A clones. Specific Hdac3 band is indicated. **(B,D)** Similar to **Figure 1B**, but using different embryo RNAseq samples (Mohammed et al., 2017) (B) or *in vitro* cell types (Anderson et al., 2017) (D). Primitive streak (PS) Extraembryonic endoderm cell states (nEnd, XEN), anterior definitive endoderm (ADE) **(C)** Similar to **Figure 1C**, but using different embryo RNAseq samples (Mohammed et al., 2017). **(E)** Similar to **Figure 1D** with a magnified view of cluster 5. **(F)** p-values of GO terms enriched in cluster 5. **(G)** Indicated PrE marker mRNA expression relative to negative control siRNA transfection of cells transfected with indicated siRNAs after 2d in SL. **(H)** *Gata6* mRNA levels relative to *WT* cells of indicated genotypes after 2d in SL. Average and standard deviation (SD) of at least two independent clones. **(I)** Similar to **Figure 1D** with a magnified view of a panel of naïve and general pluripotency, and post-implantation markers. **(J)** Anti-Hdac3 western-blot of naïve *Hdac3*^**-/-**^ G6C18 clones. Specific Hdac3 band is indicated. **(K)**Representative *Nanog*>GFP and *Gata6*::mCherry fluorescence intensity plots used for quantification in **Figure 1F**. **(L)** Similar to **Figure 1G**.

**Figure S2:**
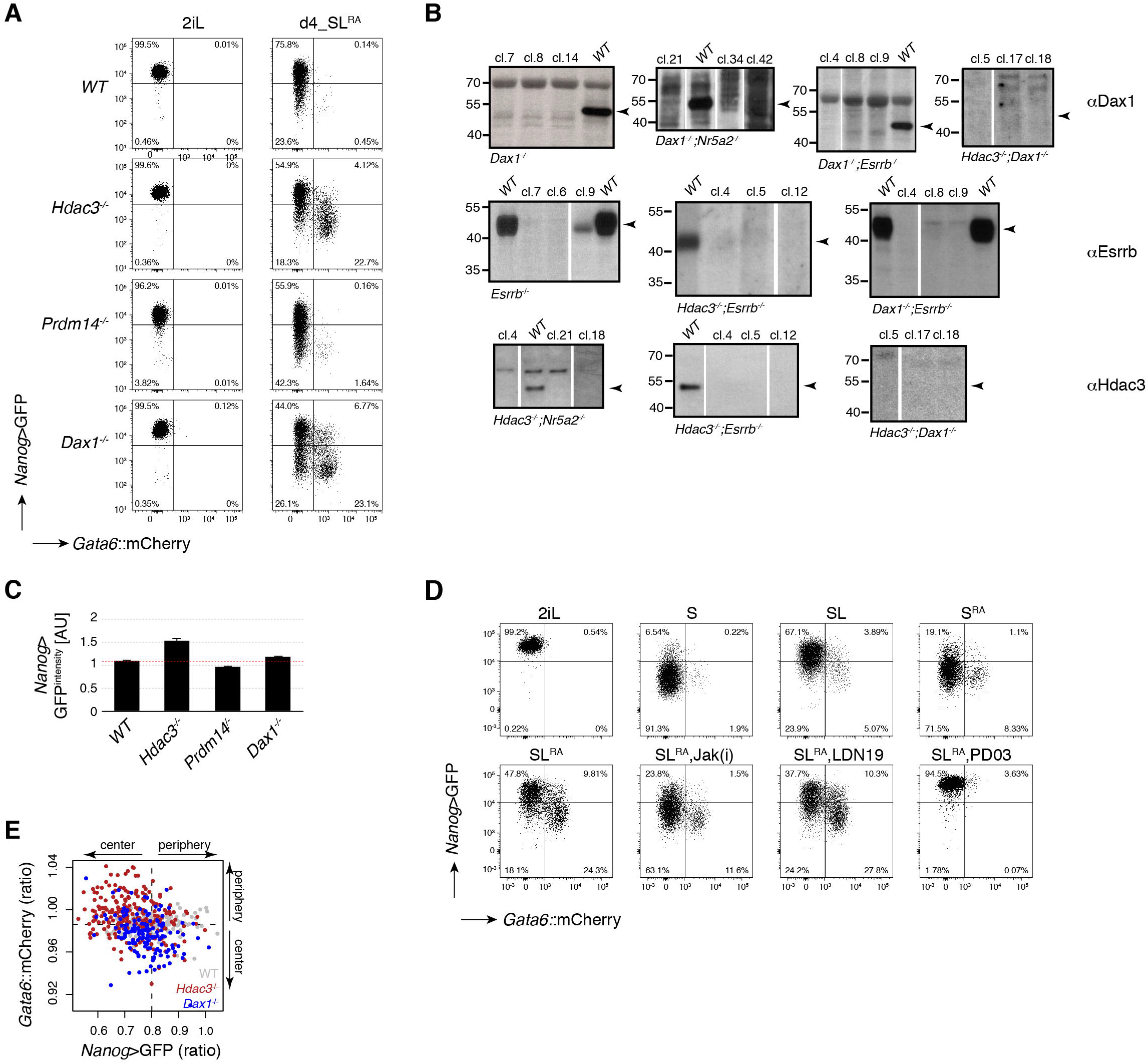
**Characterization of *Dax1*^-/-^ cells. Related to Figure 2.** **(A,D)** Representative *Nanog*>GFP and *Gata6*::mCherry intensity plots in indicated genotypes and conditions. **(B)** Anti-Dax1, -Esrrb, and -Hdac3 western blots for the genotyping of G6C18 mutant clones. Migration behavior of targeted proteins is indicated. **(C)** Quantification of Nanog>GFP geometric mean intensities in 2iL. Average and SD of at least two independent clones. **(E)** Ratios of *Nanog*>GFP and *Gata6*::mCherry intensity (periphery/center) in spheroids of indicated genotypes. Dashed lines indicate the mean in *WT* cells.

**Figure S3:**
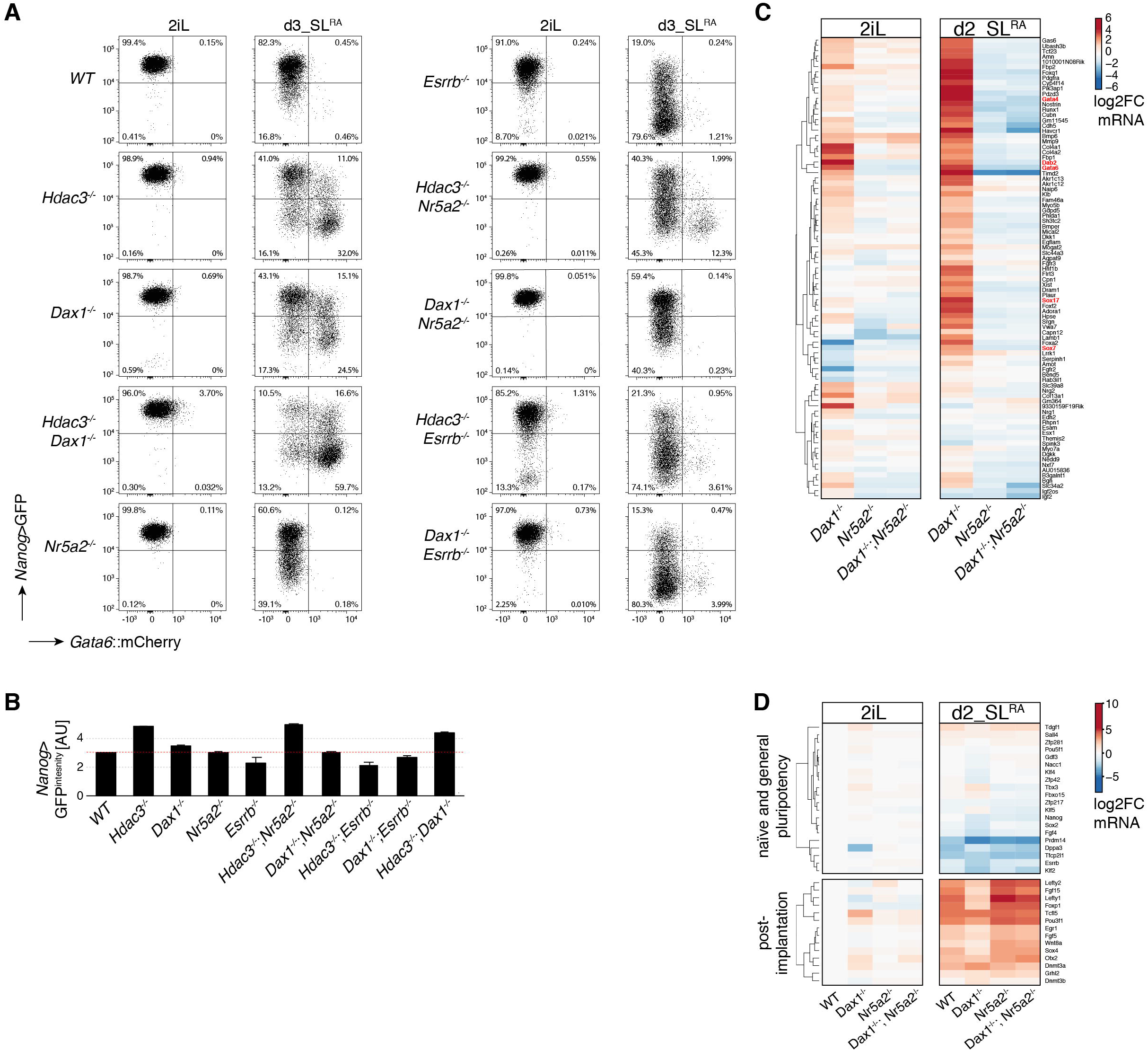
**Characterization of compound mutant cells. Related to Figure 3.** **(A)**Representative *Nanog*>GFP and *Gata6*::mCherry intensity plots of genotypes and conditions quantified in **Figure 3C**. **(B)**Geometric mean intensity of *Nanog*>GFP reporter in 2iLIF. Average and SD of three independent clones. **(C)** Similar to **Figure 3D**, but focusing on cluster 5 genes. **(D)** Similar to **Figure 1D** with a magnified view of a panel of naïve and general pluripotency, and post-implantation markers in indicated genotypes and conditions relative to naïve *WT* cells in 2iL.

**Figure S4:**
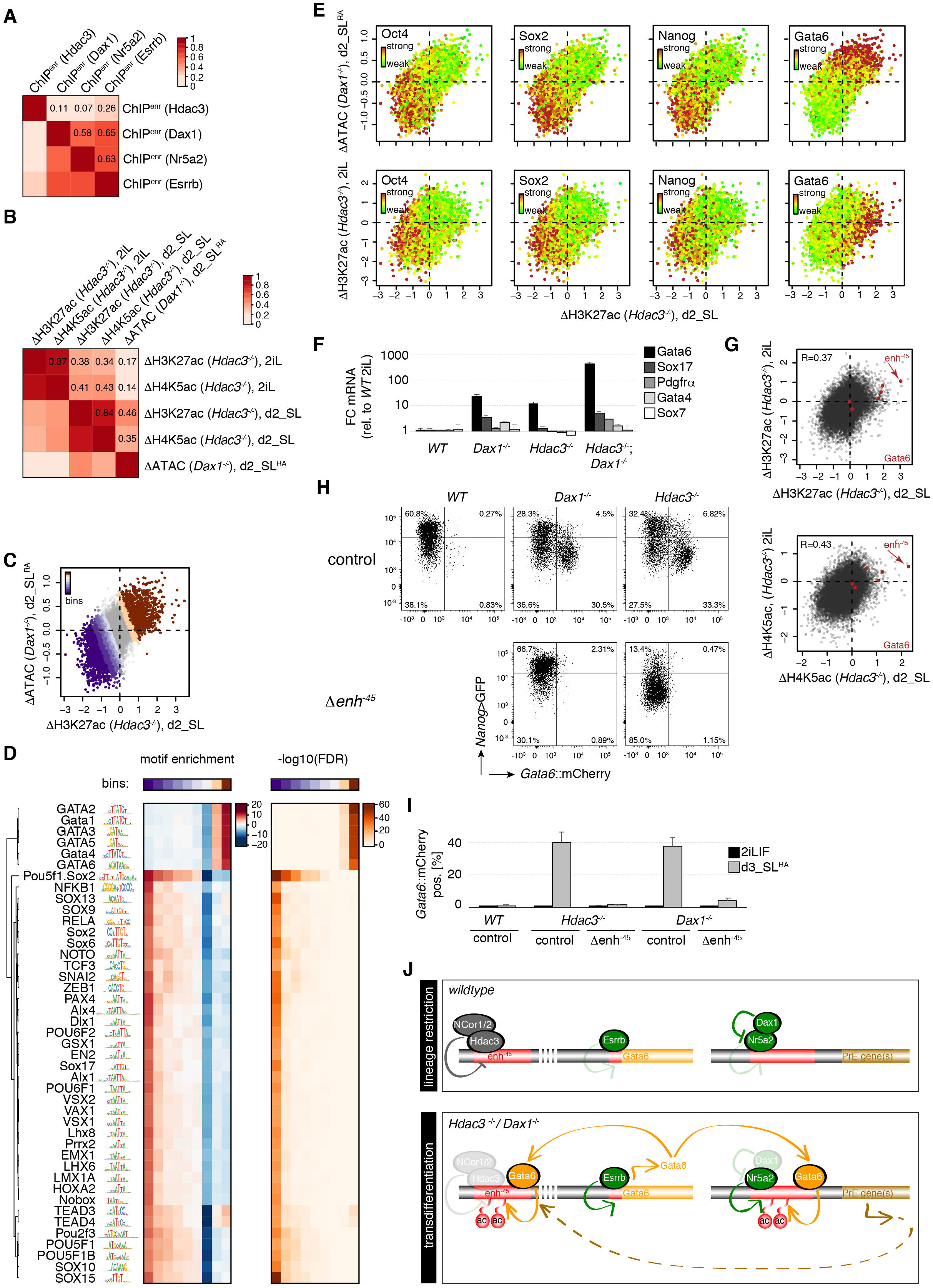
Epigenomic analysis and enh^-45^ characterization. **(A)** Pairwise Pearson correlation of Hdac3, Dax1, Nr5a2, Esrrb occupancy at regions bound by Hdac3 and/or Dax1. Individual correlation coefficients are indicated. **(B)** Pairwise Pearson correlation of changes in H3K27ac, H4K5ac or accessibility in mutant relative to *WT* cells and in indicated conditions at regions bound by Hdac3 and/or Dax1. Individual correlation coefficients are indicated. **(C)** Similar to **Figure 4B**. CRE bins according to activation and repression in *Dax1* and *Hdac3* mutants during differentiation are colored. **(D)** Heat-map of high-confidence TF motifs, enrichments and false discovery rates (FDRs) at these bins. **(E)** Similar to **Figure 4B** with individual regions colored by ChIPseq signal for indicated TFs (upper panels). The same scatterplots with log2 fold H3K27ac changes in naïve *Hdac3* mutants instead of accessibility changes in differentiating *Dax1*^**-/-**^ cells plotted on the y-axis (lower panels). **(F)** Expression changes of PrE markers in naïve ESCs of indicated genotypes relative to *WT* cells. Average and SD of three independent clones. **(G)** Scatterplots of log2 fold changes in H4K5ac and H3K27ac in *Hdac3* mutants at regions bound by Dax1 and/or Hdac3 in naïve and differentiation conditions. Regions colored in red are associated with Gata6. **(H,I)** Representative *Nanog*>GFP and *Gata6*::mCherry intensity plots (H) and quantification of *Gata6*::mCherry positive cells (I) in indicated genotypes after 3d in SL^RA^. Average and SD of at least three independent clones. **(J)** Model. In *WT* cells, Hdac3 and Dax1 form a regulatory network that inhibits lineage conversion. Hdac3/NCor1/NCor2 bind and silence enh^-45^, while Dax1 interacts with and antagonizes Nr5a2, and indirectly influences enh^-45^. Derepression of *Gata6* in *Dax1* and *Hdac3* mutants causes Gata6 to associate with Hdac3- and Dax1/Nr5a2-bound CREs, together with Esrrb forming a positive feed-forward loop for PrE transdifferentiation.

**Table S1: Oligonucleotides, siRNAs and antibodies, and sequencing information of genome-engineered clones used in this study**.

**Table S2: Data presented in Figures 1 and S1**.

**Table S3: Data presented in Figures 1, 2, S1 and S2**.

**Table S4: Data presented in Figures 4 and S4**.

